# Deletion of RSK2 kinase alleviates age-dependent hypertension

**DOI:** 10.1101/2025.03.12.642932

**Authors:** Ramon Ayon, Yves T. Wang, Jaspreet Kalra, Li Jin, Yen-Lin Chen, Renata Polanowska-Grabowska, Swapnil K. Sonkusare, Christine K. Christie, Eric M. Small, Thu H. Le, Avril V. Somlyo

## Abstract

**Background:** Hypertension prevalence increases with age, reaching over 70% of people over age 65. The underlying mechanisms are poorly understood. This study interrogates a new signaling pathway in vascular smooth muscle of aged mice driven by p90 ribosomal S6 kinase, RSK2, and its role in increasing peripheral vascular resistance and blood pressure (BP).

**Methods:** Basal BP measurements were taken at 26-29 month (812-892 day) old mice with global deletion of RSK2 (*Rsk2*^*-/-*^) prior to and following treatment with L-NAME. Cardiac function, vessel stiffness, myogenic responses, Ca^2+^events, contractility, immuno-staining, histology studies and western blotting were performed.

**Results:** Resting BP and myogenic vasoconstriction were normal in aged global *Rsk2*^*-/-*^ mice and elevated in wild type (WT) littermates. L-NAME treatment increased BP in aged *Rsk2*^*+/+*^ but not aged *Rsk2*^*-/-*^. Vessel stiffness and glycation collagen crosslinking increased in both aged *Rsk2*^*+/+*^ and *Rsk2*^*-/-*^ compared to young vessels with no remodeling or increase in collagen content, even though BP in aged *Rsk2*^*-/-*^ arterioles was normal. Increased vessel stiffness was dissociated from increased BP. Ca^2+^ transients increased and sensitivity to NO-induced relaxation decreased in aged *Rsk2*^*+/+*^ compared to young WT arterioles. IEL structures, eNOS and Hbα distribution at myoendothelial junctions were disturbed impairing vasorelaxation in aged *Rsk2*^*+/+*^ but not aged *Rsk2*^*-/-*^ arterioles.

**Conclusions:** RSK2 plays a significant role in hypertension associated with aging by downregulating prorelaxant signaling and promoting procontractile events in the vasculature, offering potential new therapeutic targets.

## Introduction

Hypertension is associated with aging with approximately 70% of elderly patients (≥65 years of age) being treated for systolic hypertension ^1^, a primary risk factor for myocardial infarction, stroke, neural injury, organ damage and cognitive impairment. ∼50% of all hypertensive patients do not respond adequately to current antihypertensive therapies ^2^. Thus, a better understanding of the mechanisms underlying this age-related increased blood pressure (BP) warrants further investigation with the potential for new therapies.

The age-related increase in BP is generally viewed to be a hallmark of aging blood vessels and is ascribed to endothelial dysfunction and vascular remodeling with stiffening and fibrosis of the vessel wall ^3^. The etiology of the initial rise in BP is unresolved ^4^. There are contrary findings to this generally accepted view. For example, elevated BP in 1-year-old mice has been reported to occur without changes in vessel stiffness or renal or endocrine abnormalities ^5,6^. Likewise, SM cell mineralocorticoid receptors or G-protein signaling contribute to vascular tone and BP regulation independent of vascular structural changes and renal defects in sodium handling ^5-7^. Our focus on the role of the vasculature in age related hypertension stems from our novel finding that resting BP is lower in young ^8^ and in aged transgenic mice lacking p90 ribosomal S6 kinase (RSK2) than in WT littermates. We have shown in young mice, that RSK2 contributes to BP, myogenic tone and blood flow and is responsible for ∼25% of the maximal contractile force in resistance arteries ^8^. These cardiovascular effects reflect the ability of RSK2 to directly phosphorylate SM myosin regulatory light chain (RLC_20_) and to phosphorylate an activating site in the Na^+^/H^+^ exchanger (NHE-1), resulting in intracellular alkalinization, an increase in cytosolic Ca^2+^ resulting in vasoconstriction ^8^. Increased activity of RSK2^Ser227-P^ and NHE-1^Ser703-P^ associated with hypertension in rat and human SM cells has been reported, but why that lead to increased BP was not understood ^9-11^. RSK2^Ser227-P^ has also been found in senescent human renal arteries ^12^ and to a lesser extent in mesenteric arteries but BP or RSK2 deletion was not studied.

Age-induced endothelial dysfunction of the microcirculation with impairment of the NO dilatory pathway through decreased production or increased degradation of eNOS and NO and increased reactive oxygen species results in increased BP^13,14^. In resistance arteries the signaling hubs between the NO generating endothelial cells and SM occur at holes in the internal elastic lamina (IEL) where the endothelial cell (EC) projections contact the underlying SM cells forming myoendothelial junctions (MEJs). RSK2 may alter these structures and distribution of the associated NO, eNOS and the NO scavenger, Hbα, in aged resistance vessels ^15,16^ that could regulate NO-induced relaxation and blood pressure.

The goal of this study was to test our hypothesis that RSK2 signaling, through regulation of procontractile and prorelaxant mechanisms and through structural changes and functional changes at the IEL/MEJ and remodeling in resistance arteries contributes to hypertension associated with aging.

## Methods

A detailed description of all methods can be found in Supplemental Material

## Results

### RSK2 deletion protects against development of hypertension and response to eNOS inhibitor LNAME with aging

RSK2 protein expression was similar in aged (78-87 week old) WT compared to young (10-15 week old) mice and was undetectable in aged global *Rsk2*^*-/-*^ mice (Fig. 1A). Basal systolic BP (SBP) was elevated in aged *Rsk2*^*+/+*^ mice but was normal in aged *Rsk2*^*-/-*^ mice (123 ± 10 vs 99 ± 12 mm Hg (Fig 1B). As the aged mice were fragile and at risk of dying from surgical placement of radiotelemetry transmitters, BP was measured using tail cuff manometry. In our hands, BP measurements are reproduceable when comparing the 2 approaches (3). Shown in Table 1 a similar difference in BP between WT and *Rsk2*^*-/-*^ mice was found using both techniques, with tail cuff measurements lower than telemetric. In the L-NAME model of hypertension, SBP increased in WT aged mice by 14 mm Hg to 137 ± 2 mm Hg, while, surprisingly, the aged *Rsk2*^*-/-*^ mice were refractory to L-NAME treatments, with no induced hypertension, SBP 97 ± 8 mmHg (Fig. 1B). Heart rate at baseline did not differ between WT and *Rsk2*^*-/-*^ mice while L-NAME induced bradycardia in both groups (Fig. 1C), consistent with previous reports in rats ^17^. Echocardiographic studies were performed at baseline and no significant differences found in the left ventricular parameters, including ejection fractions, between WT and *Rsk2*^*-/-*^ aged mice, Table 2. Age related mortality in this cohort of mice was greater in aged WT (7 of 11, 63.6% over 2-5 months) compared to aged *Rsk2*^*-/-*^ mice (1 of 8, 12.5% over 5 months). Thus, the ability of aged *Rsk2*^*-/-*^ mice to maintain a normal resting BP even when challenged with L-NAME implicates a significant role for RSK2 signaling in the regulation of BP and hypertension associated with aging.

**Table 1:** SBP under basal conditions in young (10-15 weeks) and aged (88-91 weeks) global *Rsk2*^*-/-*^ and WT littermates as well as *Rsk2*_*SM-/-*_ using radiotelemetry or tail-cuff measurements. Young *Rsk2*_*-/-*_ mice, in which RSK2 expression is specifically reduced by ∼70% in SM displayed significantly reduced SBP similar to measurements in the global *Rsk2*^*-/-*^ indicating that the effects of RSK2 on SBP are primarily on the vasculature.

|  | SBP Telemetry<br>x ± SEM |  | SBP Tail Cuff<br>x ± SEM |  |
| --- | --- | --- | --- | --- |
|  | RSK2 global<br>KO<br>(Young) | RSK2<br>conditional KO<br>(Young) | RSK2 global<br>KO<br>(Young) | RSK2 global<br>KO<br>(Aged) |
| WT | 124.4 ± 0.7 | 120.8 ± 4.0 | 108.1 ± 1.7 | 122.8 ± 10.0 |
| KO | 112.7 ± 0.7 | 109.5 ± 7.1 | 96.1 ± 2.3 | 98.5 ± 12 |
| Diff | 13 | 11 | 12 | 24.3 |
| n | 7/3 | 4/10 | 15/9 | 9/7 |
| p (two-tailed) | 0.03 | 0.004 | 0.0002 | 0.001 |

**Table 2:**
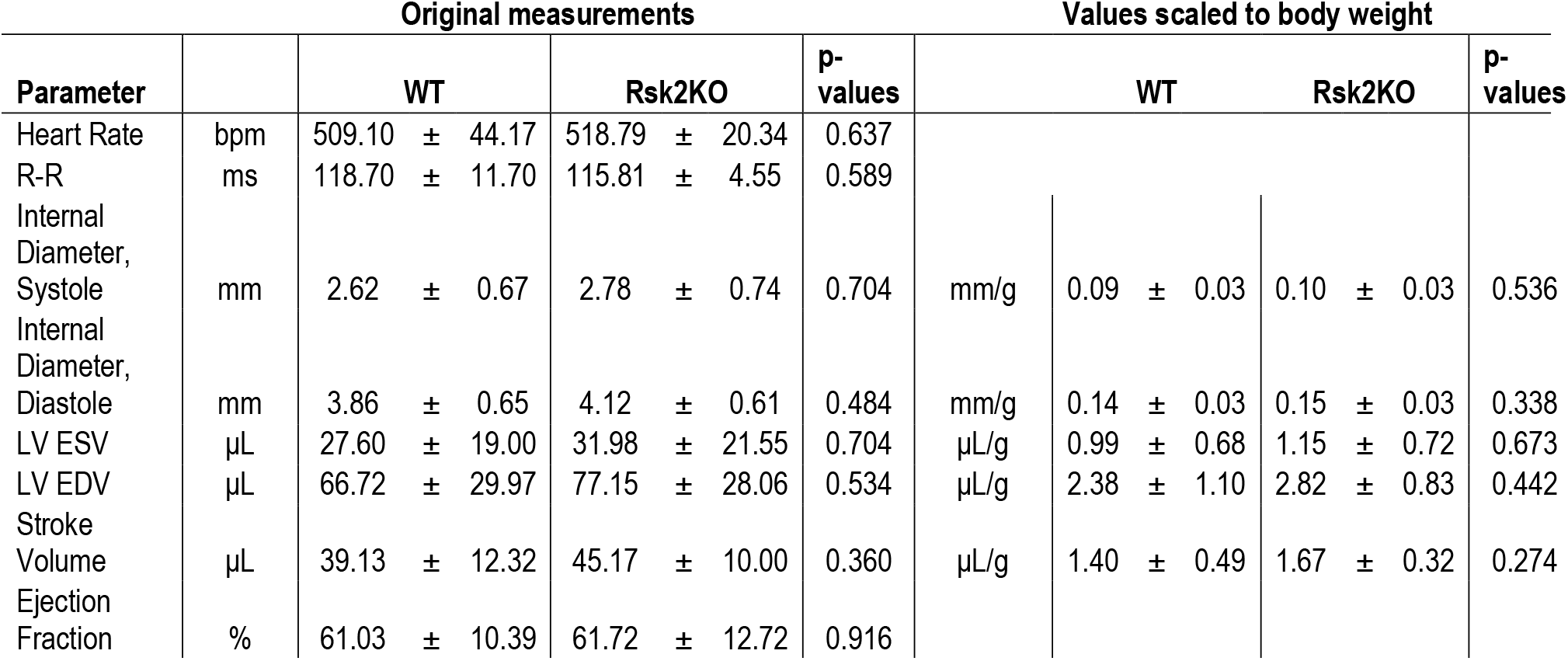
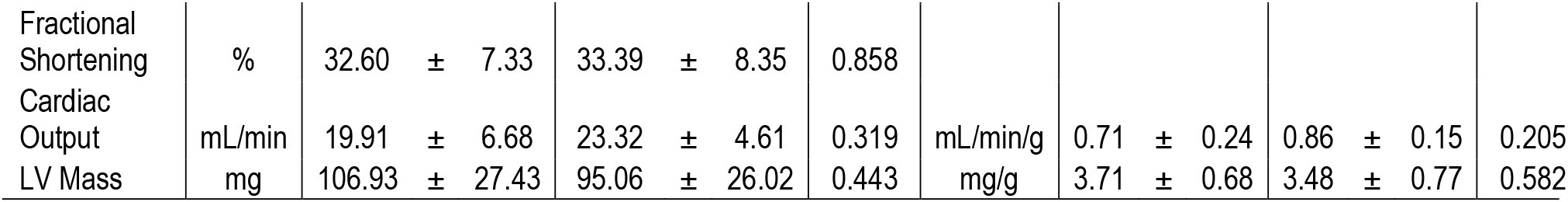
Cardiac functions measured by MRI in aged *Rsk2*^*+/+*^ and aged *Rsk2*^*-/-*^ littermates. LV Mass – Left Ventricular Mass, LV ESV – Left Ventricular End Systolic Volume, LV EDV - Left Ventricular End Diastolic Volume, R-R – R-R wave ECG interval.

**Fig. 1:**
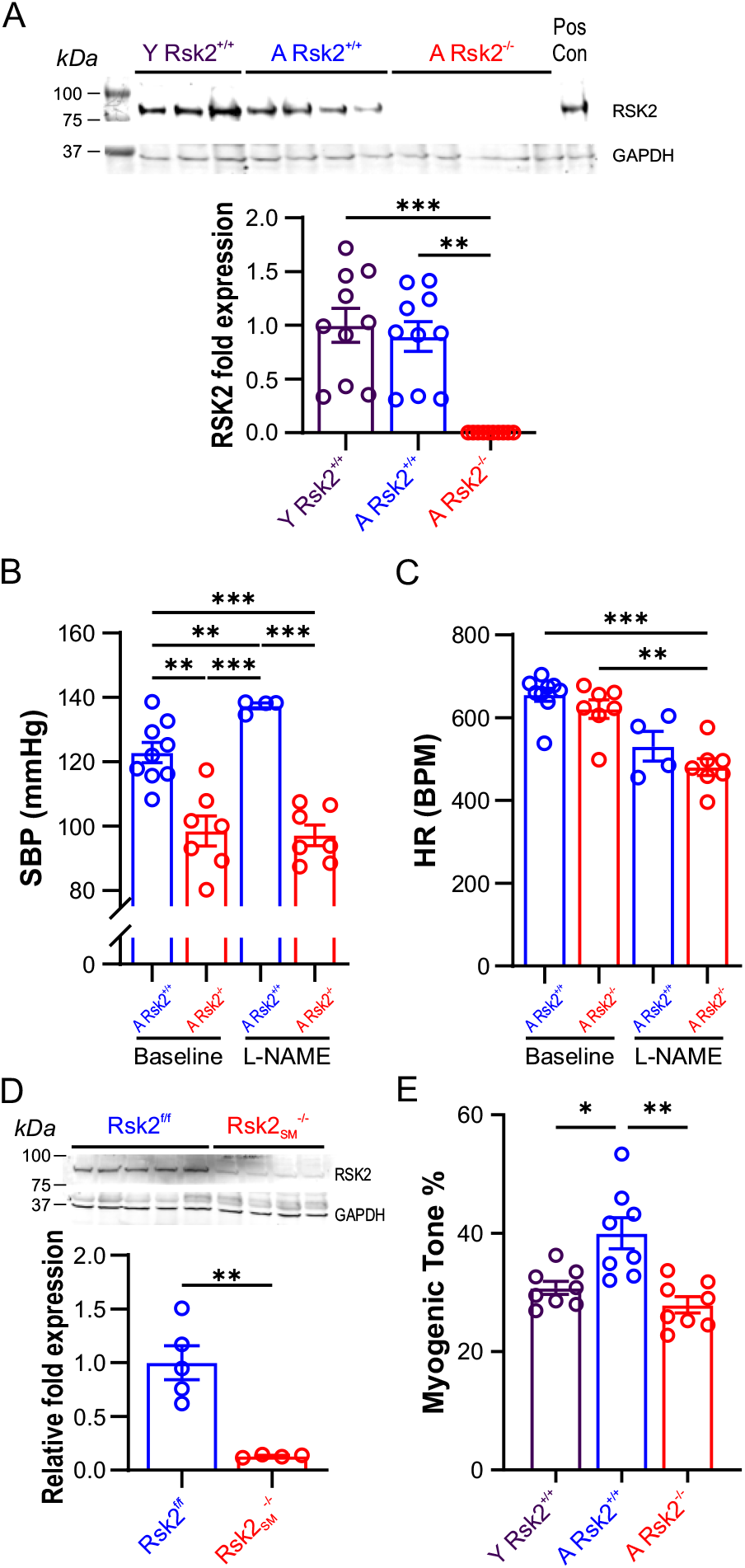
Deletion of RSK2 lowers SBP, is resistant to L-NAME treatment, and decreases myogenic tone associated with aging. (A) Representative western blot and bar graph showing RSK2 protein expression in mesenteric arteries from young (Y) RSK2^+/+^, aged (A) RSK2^+/+^, and A RSK2^-/-^ mice. Liver homogenate = a positive control, and GAPDH a loading control. A significant reduction in RSK2 expression was detected in A RSK2^-/-^ mice but not between Y and A RSK2^+/+^ mice (n = 10 per group). (B) SBP was significantly lower in A RSK2^-/-^ mice compared to A RSK2^+/+^ mice at baseline and following L-NAME treatment (n = 7-9 mice per group). (C) Heart rates were NS between A RSK2^-/-^ and A RSK2^+/+^ mice at baseline or following L-NAME treatment. However, L-NAME induced bradycardia (n = 7-9 mice per group). (D) RSK2 expression in SM-specific RSK2 knockout (RSK2_SM_^-/-^) mice was reduced by approximately 87% following tamoxifen induction (n = 4-5 per group). (E) Myogenic responses of mesenteric resistance arteries from Y RSK2^+/+^ (n = 4 mice, 8 arteries), A RSK2^+/+^ (n = 4 mice, 8 arteries), and A RSK2^-/-^ (n = 4 mice, 8 arteries) at 80 mmHg. Myogenic tone was significantly greater in A RSK2^+/+^ mice compared to Y RSK2^+/+^, but no difference was observed between Y RSK2^+/+^ and A RSK2^-/-^ mice. (\***P** < 0.05, \*\***P** < 0.01, \*\*\***P** < 0.001).

### Smooth muscle (SM) specific deletion of Rsk2 results in lower BP

*Rsk2Myh11-Cre*^*+*^*(Rsk2* ^*-/-*^) mice specifically targeted to SM were compared to *Rsk2*^*fl/fl*^ *(Rsk2* _*SM*_^*+/+*^) littermates. Basal BP measured by radiotelemetry following oral tamoxifen in 19-22 week young mice to suppress Rsk2 expression specifically in SM (*Rsk2* _*SM*_^*-/-*^) by ∼ 80% (Fig. 1D) was significantly lower than *Rsk2* _*SM*_^*+/+*^ littermates, similar to our previous findings on global *Rsk2* _*SM*_^*-/-*^ mice (21-65 week old mice (3), (Table 2). These findings on the *RSK2 knockout mice* demonstrate a vascular basis for the role of RSK2 in basal BP control. However, for studies of the effects of RSK2 on vascular aging we have used the global *Rsk2*^*-/-*^ mice rather than the *Rsk2* _*SM*_ ^*-/-*^, because of the lack of an established protocol for multiple tamoxifen administrations and monitoring RSK2 quantitation needed over 78-87 weeks to generate a population of RSK2 deficient aged mice.

*Myogenic vasoconstriction is increased in resistance arteries in aged Rsk2*^*+/+*^ *mice but not in aged Rsk2*^*-/-*^ *mice* Vasoconstriction in response to increases in intraluminal pressure in resistance arteries is known as a myogenic response and is important for the autoregulation of blood flow for maintenance of tissue oxygenation. 80 mm Hg intraluminal pressure applied to 3^rd^ and 4^th^ order mesenteric arteries mounted on a pressure myograph led to a 39.98 ± 2.62% decrease in lumen diameter in 88 week old WT mice compared with 27.21 ± 3.30% in aged matched *Rsk2*^*-/-*^ mice (Fig. 1E). Thus, the absence of RSK2 in aged resistance arteries significantly decreases myogenic vasoconstriction.

### RSK2 signaling contributes to enhanced calcium signaling in arterioles from aged mice

Ca^2+^ events in SMCs were measured in pressurized mesenteric arteries (89 mm Hg) loaded with the Ca^2+^ indicator dye fluo-4 (Fig. 2). The frequency of spontaneous Ca^2+^ transients was greater in aged (78-87 week old) compared to young (10-15 week old) mice (0.30 ± 0.06 Hz vs. 0.12 ± 0.03 Hz), (Fig. 2 B,C) and was not altered by treatment with the RSK inhibitor, LJH685. The mean Ca^2+^ amplitude tended to be upregulated in mesenteric arteries from aged compared to young mice (1.36 ± 0.02 F/Fo vs 1.27 ± 0.02 F/Fo), while LJH685 treatment significantly decreased the amplitude of Ca^2+^ events in both young (1.18 ± 0.01 F/Fo) and aged (1.21 ± 0.01 F/F0) mice (Fig 2C). Therefore, the predominant effect of RSK2 on the increased Ca^2+^ signaling associated with aging is on the amplitude of the Ca^2+^ events.

**Fig. 2:**
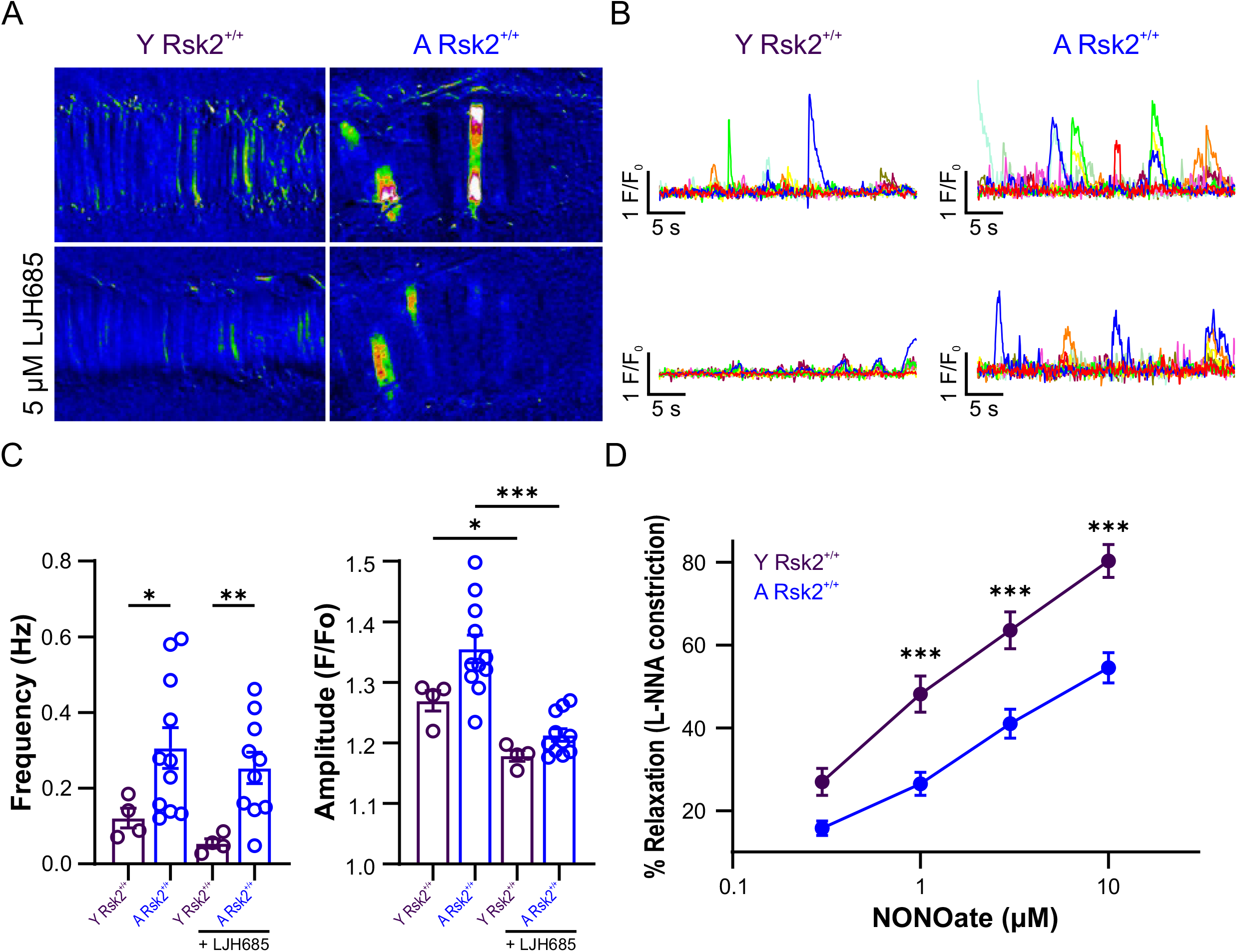
RSK2 signaling contributes to enhanced calcium signaling in arterioles from aged mice and aging reduces NO-induced vasodilation. (A) Topographic color map of Ca^2^+signaling recorded in mesenteric arteries (MAs) loaded with Fluo-4 AM (10 µM) from young (13 weeks) and aged (88 weeks) C57BL/6J mice in the presence and absence of the RSK inhibitor LJH685 (5 µM). (B) Representative Ca^2^+traces from 10 regions of interest showing spontaneous Ca^2^+activity at baseline and in the presence of LJH685 (5 µM) in young and aged MAs. (C) Summary of Ca^2^+frequency (Hz) and amplitude (F/F_0_) changes at baseline and following treatment with LJH685 (5 µM) in young WT (n = 4 mice) and aged WT (n = 11 mice). (D) Sensitivity to NO was assessed by applying the NO donor spermine NONOate (0.3–10 μM) in the presence of the NOS inhibitor L-NNA (100 μM) to pressurized (80 mmHg) MAs from young WT (n = 12) and aged WT (n = 12) mice. Aging significantly reduced NO-induced vasodilation. Data are presented as mean ± SEM. (\***P** < 0.05, \*\***P** < 0.01, \*\*\***P** < 0.001).

### NO-induced relaxation was attenuated with aging of arterial SM

Relaxation of pressurized (80 mm Hg) mesenteric arteries preconstricted with L-NNA to inhibit production of endogenous NO by NOS was measured. Subsequent relaxation in response to increasing concentrations of the NO donor spermine NONOate was measured in arteries from WT young and aged mice. Relaxation responses were significantly attenuated (p<0.05) at both 1, 3 and 10 µM NONOate in the aged compared to the young arteries (Fig. 2D).

### Aging alters the structure of the internal elastic membrane and the distribution of eNOS and Hbα at myoendothelial junctions

Serendipitously, during immuno-histology studies, we observed in the 12 and 24 month old WT, compared to the young WT and the aged *Rsk2*^*-/-*^ mesenteric arteries viewed *en face*, a significant decrease in number and an increase in size of the holes in the IEL (Fig.3AB). Myoendothelial junctions occur at the holes. Immunolabeling revealed a change in localization of eNOS that was absent or diffuse at the MEJs/IEL sites in aged *Rsk2*^*+/+*^ arteries but pinpoint in the center of the holes in both young animals and in aged *Rsk2*^*-/-*^ mice (Fig.3A). The PLA assay used to detect eNOS/Hbα complexes showed a significant decrease in puncta in the aged *Rsk2*^*-/-*^ compared to the aged *Rsk2*^*+/+*^ (Fig. 3D)

**Fig. 3:**
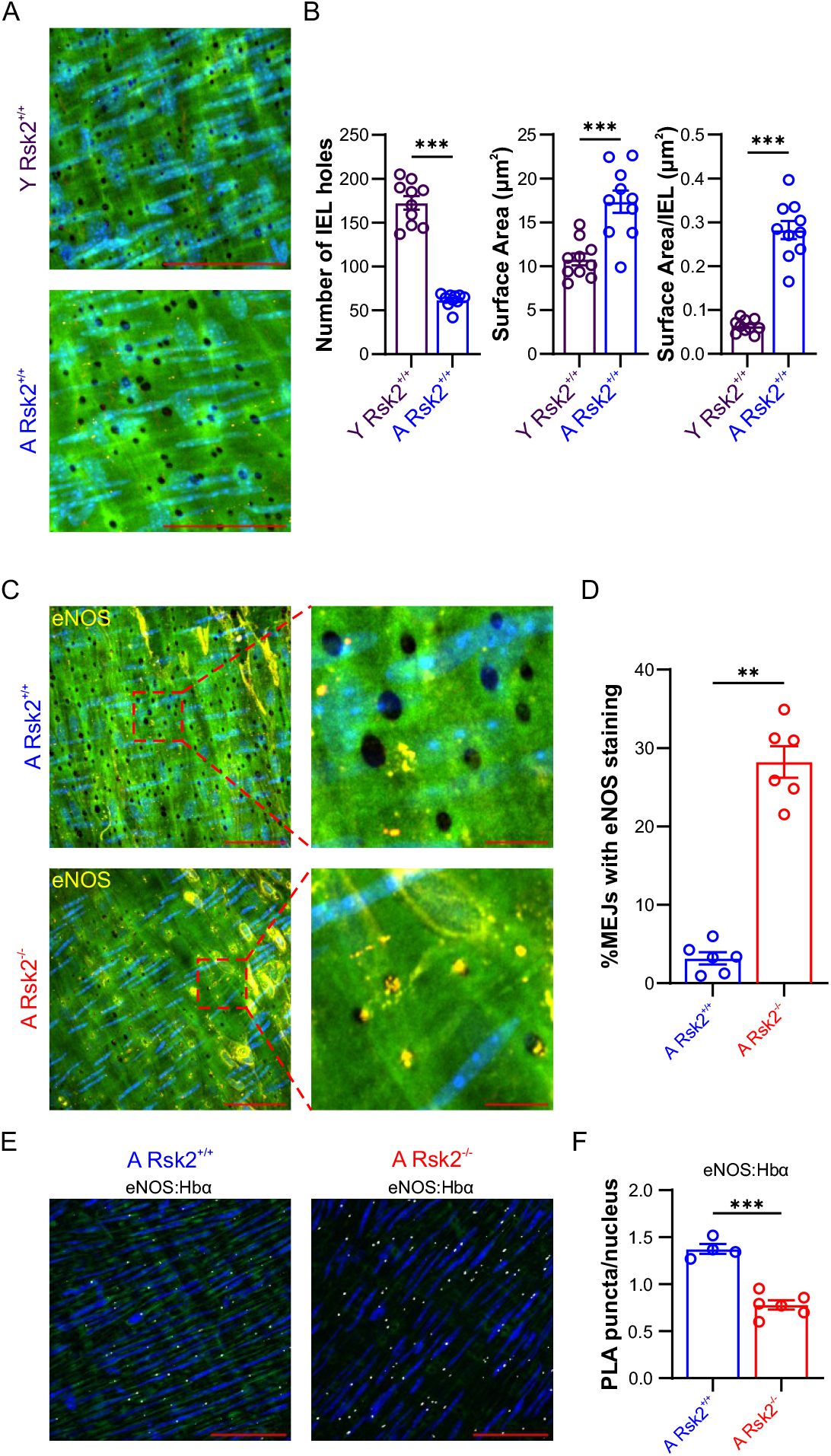
Aging alters the structure of the internal elastic membrane (IEL) and the distribution of eNOS and Hbα at myoendothelial junctions. (A) Representative immunofluorescence images of *en face* second-order MAs from Y Rsk2^+/+^ and A RSK2^+/+^ mice, showing the organization of the internal elastic lamina (IEL). (B) Quantification of IEL characteristics: (left) total number of IEL holes within the field of view, (middle) total hole surface area, and (right) surface area normalized to the total number of IEL holes within the region of interest. A Rsk2^+/+^ MAs exhibited a significantly lower number of IEL holes but a significantly larger IEL surface area compared to Y Rsk2^+/+^ mice (n = 10). (C) Representative merged immunofluorescence images of *en face* preparations of second-order MAs from A Rsk2^+/+^ and A Rsk2^-/-^ mice, showing IEL autofluorescence (green) and eNOS immunofluorescence (yellow). Black holes within the IEL represent MEJs. (D) Quantification of the percentage of eNOS immunofluorescence localized at MEPs (**n** = 6). (E) Representative proximity ligation assay (PLA) merged images showing nuclei (blue) and white puncta indicating eNOS:Hbα co-localization in *en face* preparations of MAs from A Rsk2^+/+^ and A Rsk2^-/-^ mice. (F) Quantification of eNOS:Hbα PLA puncta in MAs, normalized to nuclei (**n** = 4-6). (\*\*\***P** < 0.001).

### Aging did not result in significant vascular remodeling or increased collagen content of mesenteric arterioles and iliac arteries

Lumen diameter was greater in the aged *Rsk2*^*+/+*^ and aged *Rsk2*^*-/-*^ mesenteric and iliac arteries compared to young arteries (Fig. 4 A-C) as shown in Masson’s trichrome-stained sections. Wall thickness was slightly increased in the aged mesenteric arteries relative to young ones (Fig. 4A,B), likely reflecting overall vessel size in older mice. However, no differences were observed when wall thickness was normalized to total wall area (Fig. 4B,D). Collagen content was evaluated using polarized light of picrosirius red-stained cross sections of mesenteric arterioles and iliac arteries (Fig. 4E,F). There were NS differences between any groups.

**Fig. 4:**
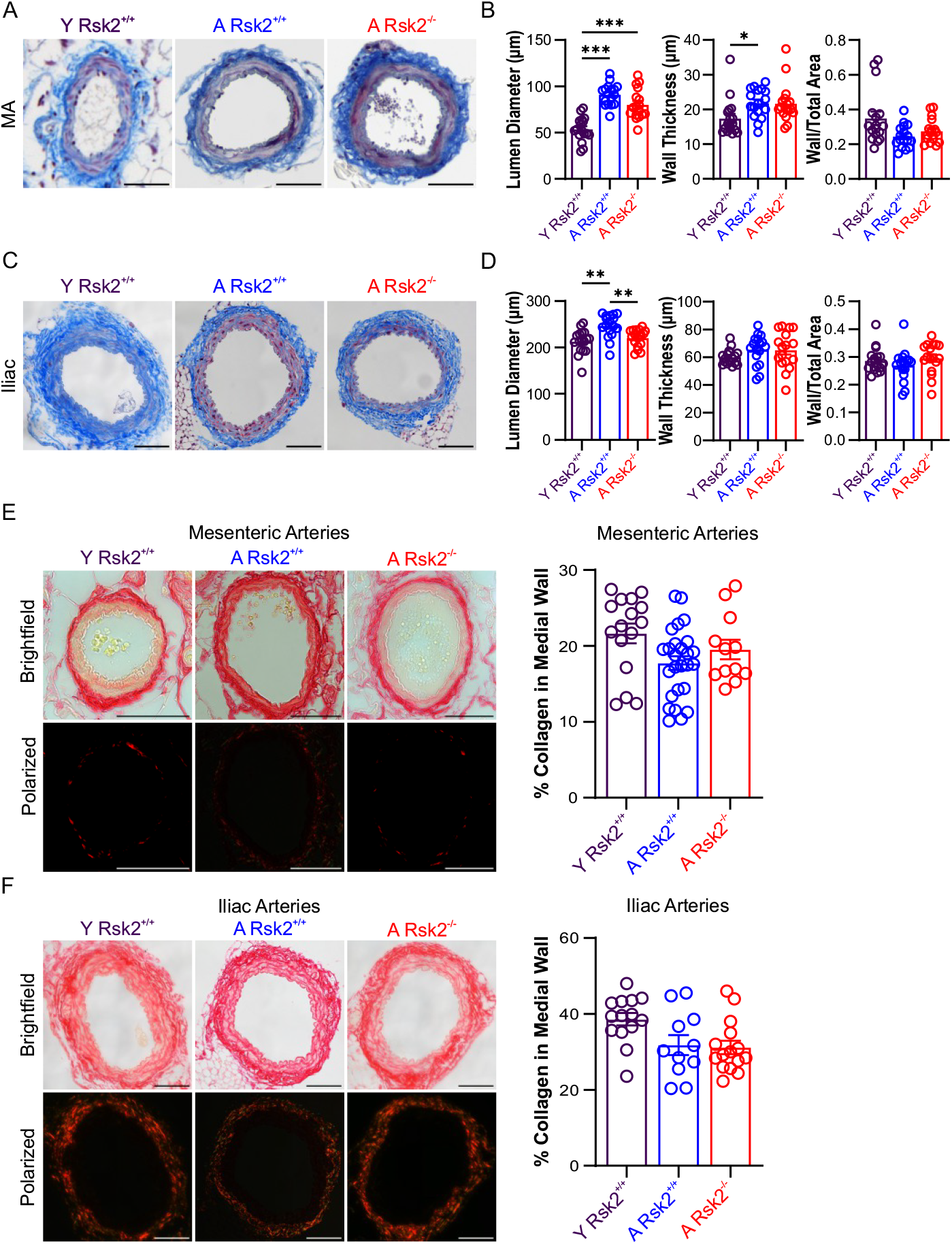
Aging did not significantly result in vascular remodeling and increased collagen content of mesenteric arterioles and iliac arteries. (A-D) Representative images of mesenteric arterioles (MA) and iliac arteries stained with Masson’s trichrome, in young and aged mice. Lumen diameters were significantly larger in aged mesenteric arterioles and iliac arteries. Wall thickness did not differ in iliac arteries between groups; however, in A Rsk2^+/+^ MAs, wall thickness was significantly greater compared to Y Rsk2^+/+^ (P < 0.05). The ratio of wall area to total area between 20-month-old Rsk2^+/+^ (n = 17) and Rsk2^-/-^ (n = 17) mice did not differ, suggesting that differences in cross-sectional dimensions reflect the overall larger size of aged mice compared to young mice (Fig. 4E,F). Aging did not increase vascular collagen content, as assessed by polarized light microscopy of picrosirius red-stained cross-sections of MAs and iliac arteries (**n** = 11-26).

*Mechanical stiffness of resistance arteries was increased in both aged Rsk2*^*+/+*^ *and Rsk2*^*-/-*^ *compared to young WT*

The mechanical properties of arterioles mounted in a pressure myograph were determined under Ca^2+^-free conditions to eliminate contractile mechanisms. Arterial stiffness was calculated from passive-pressure diameters taken at 10, 20, 40, 60, 80, 100 and 120 mm Hg (Fig. 5 A,B,C). The stress/strain curves of aged *Rsk2*^*+/+*^ and *Rsk2*^*-/-*^ arteries were superimposable and left-shifted when compared to the young arteries (Fig. 5C). An increase in the stress strain relationship stiffness coefficient (ß) was observed in aged *Rsk2*^*+/+*^ (10.39 ± 1.60) and aged *Rsk2*^*-/-*^ (9.86 ± 1.08) when compared to young arteries (5.63 ± 0.31) p = 0.79 (Fig. 5C insert). Thus, deletion of RSK2 has not altered the structural changes in the aged vessel wall responsible for the increased stiffness.

**Fig. 5:**
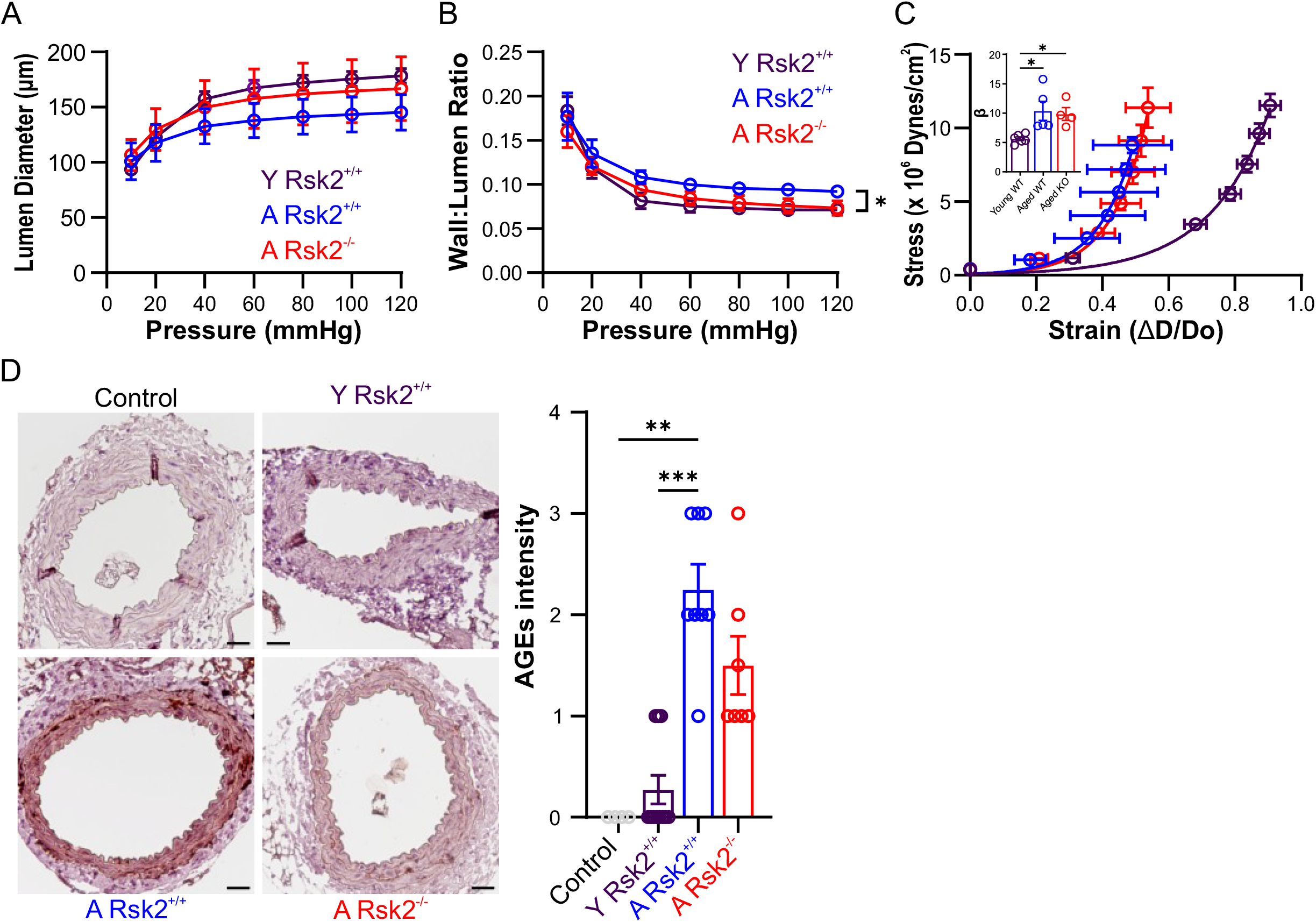
Mechanical stiffness and AGEs of resistance arteries were increased in both aged RSK2^+/+^ and Rsk2^-/-^ compared to young WT. (A-C) Comparison of structural and mechanical parameters in MAs from young and aged Rsk2^+/+^ and aged Rsk2^-/-^ mice. (A) Intraluminal diameter measurements of fully dilated MAs from A Rsk2^+/+^ (n = 5), A Rsk2^-/-^ (n = 4), and Y Rsk2^+/+^ (n = 6). (B) Wall-to-lumen ratio in MAs across all groups. (C) Stress-strain curves of fully dilated MAs. The stiffness coefficient (β) from the stress-strain relationship (inset) was significantly increased in both aged Rsk2^+/+^ and aged Rsk2^-/-^ mice compared to young Rsk2^+/+^ mice, with NS difference between the aged groups. (D) Advanced glycation end-product (AGE) crosslinking in iliac artery sections was significantly greater in aged Rsk2^+/+^ and aged Rsk2^-/-^ mice compared to young Rsk2^+/+^. Secondary antibody-only reactions served as negative controls. Vessel media staining was blindly scored using the following criteria: 0 = grey/blue (no staining), 1 = light brown staining, 2 = light brown staining with some darker deposits, 3 = light brown staining with many darker deposits. Sample sizes: n = 4-6, Y Rsk2^+/+^: 10–15 weeks, A Rsk2^+/+^: 88– 91 weeks, A Rsk2^-/-^: 88–90 weeks.

### Advance glycation products (AGEs) were increased in both aged Rsk2^+/+^ and Rsk2^-/-^ compared to young WT arteries

Detection of AGEs using immunohistochemistry was present in all aged *Rsk2*^*+/+*^ and *Rsk2*^*-/-*^ sections of iliac arteries, being most pronounced in the aged *Rsk2*^*+/+*^ samples (Fig. 5D). AGE signals were detected in two of the four arteries from young animals (2-3 months old).

### Interstitial and perivascular cardiac fibrosis in aged compared to young mice

Hypertension is a known risk factor for heart failure, which could contribute to the differences in BP in the aged *Rsk2*^*+/+*^ and *Rsk2*^*-/-*^ mice. Therefore, we evaluated evidence for cardiac fibrosis in mid-level sections of ventricles from young and aged *Rsk2*^*+/+*^ and *Rsk2*^*-/-*^ hearts using Picrosirus red staining (Suppl. Data Fig.1). Aged *Rsk2*^*+/+*^ and *Rsk2*^*-/-*^ hearts imaged with polarized light had small but significantly greater collagen content than young hearts (Suppl. Data, Fig.1E). The normal endomysium surrounding individual myocytes gave very weak variable staining in the left and right ventricles in sections of the young hearts and was significantly greater but still weak in the aged *Rsk2*^*+/+*^ and *Rsk2*^*-/-*^ samples, as was perivascular fibrosis. There was no difference in fibrosis between aged *Rsk2*^*+/+*^ and aged *Rsk2*^*-/-*^ hearts despite a difference in BP.

## Discussion

We have shown that MLCK*-*deficient mouse embryos develop to term and embryonic blood vessels contract and that RSK2 is an ancillary kinase(s) supporting contraction in the absence of MLCK ^18^. We now show that RSK2 signaling plays a significant role in hypertension associated with aging. This is based on our finding that aged (18-20 month old) *Rsk2*^*-/-*^ mice, unlike aged *Rsk2*^*+/+*^ mice, had normal BP, and remarkably, were resistant to L-NAME induced hypertension.

How does RSK2 contribute to the pathogenesis of vascular aging and hypertension? Procontractile signaling pathways, such as Endothelin-1 and SM cell mineralocorticoid receptors are known to be up regulated (5,7) as well as sympathetic nerve stimulation, alpha-adrenergic vasoconstriction ^19^, increased responsiveness to the thromboxane analogue, U46619 ^12^ and the renin-angiotensin system AngII ^3^. Furthermore, endothelin-1, AngII, U46619 and phenylephrine induce activatory phosphorylation of RSK2 in resistance arteries (see Fig. 3 in (8)). AngII stimulation of RSK2 in VSM cells inducing activation of the Na^+^/H^+^ exchanger, NHE-1^20^. Myogenic vasoconstriction also increases with aging ^21^ and was suppressed by about 25% in the aged *Rsk2*^*-/-*^ arterioles. Increased Ca^2+^ events are downstream of the above procontractile stimuli in aged arterioles, resulting in increased basal tone and BP. The increased Ca^2+^ reflects the RSK2/NHE-1 mediated alkalinization^8^. Many Ca^2+^ channels are pH sensitive, as are IP_3_- and Ca^2+^-induced Ca^2+^ release ^22^, contributing to increased cytosolic Ca^2+^ and contractility. We suggest that the surprising finding that L-NAME administration does not increase BP in aged *Rsk2*^*-/-*^ mice is because RSK2 is downstream of the above up-regulated procontractile stimuli, thus shifting the balance from procontractile in favor of prorelaxant mechanisms in the aged *Rsk2*^*-/-*^ arterioles (Fig. 6). Prorelaxant mechanisms would be enhanced in aged *Rsk2*^*-/-*^ compared to aged *Rsk2*^*+/+*^ mice due to the increased sensitivity to NO and preserved normal MEJ structure and eNOS distribution resulting in a condition favoring normal BP. Other vasodilatory mechanisms at MEJ, such as membrane hyperpolarization and TRPV4 channels may also contribute but their regulation by RSK2 has not been investigated.

**Fig. 6:**
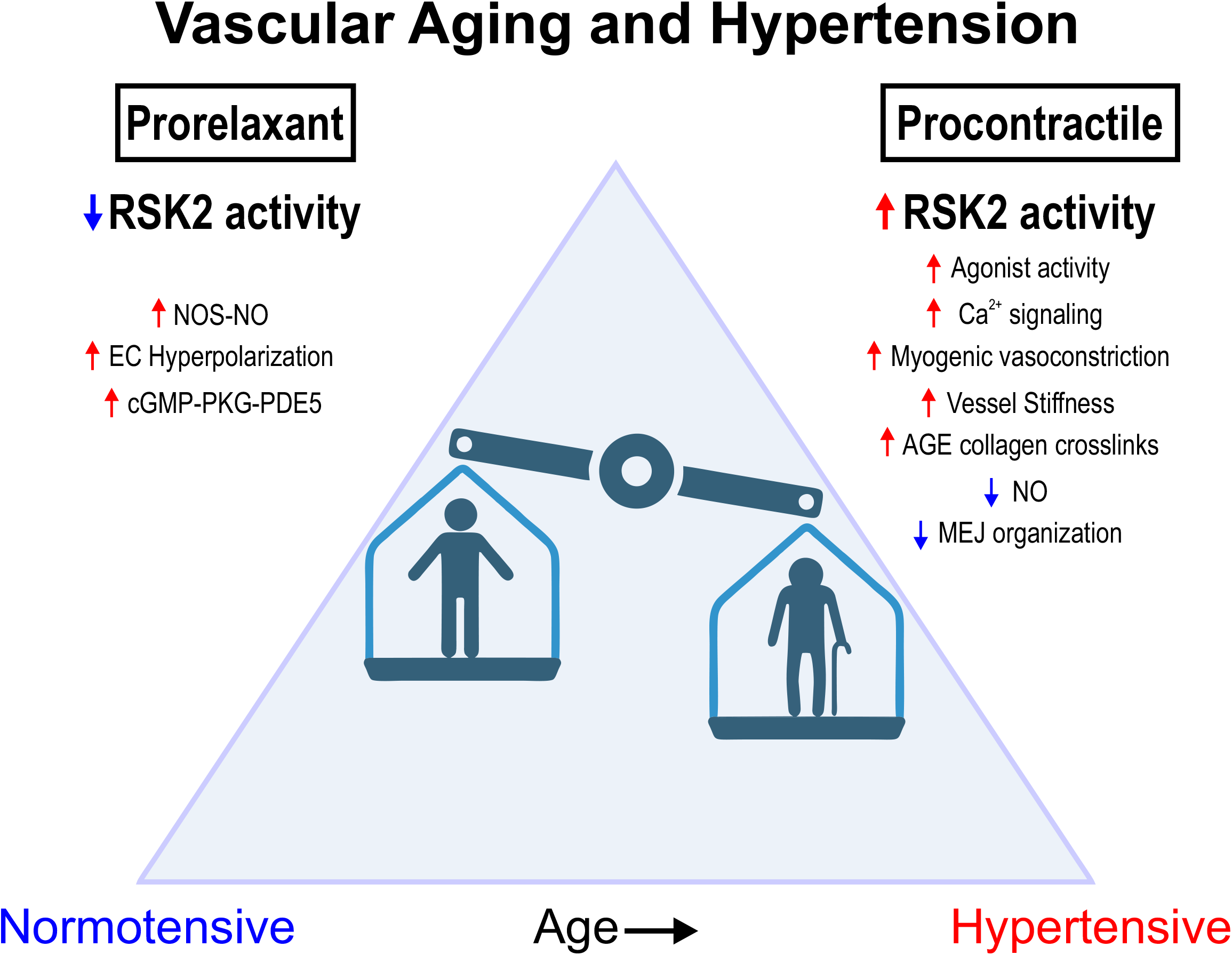
Vascular aging and hypertension scheme: RSK2 shifts the balance of prorelaxant to procontractile mechanisms. In young arteries the balance between prorelaxant and procontractile mechanisms account for resting basal tone and normal BP. With aging, up regulation of RSK2 mediated procontractile and decreased prorelaxant mechanisms tip the balance to increased BP and hypertension.

ENOS localizes to the MEJ with Hbα, known to scavenge NO and regulate NO diffusion and vasodilation ^15,23^. Endothelium-dependent vasodilation is known to be impaired in the aging rodent and human vasculature ^24-26^. Our findings of RSK2 dependence of changes in the IEL holes, the distribution of eNOS and the change in eNOS/Hbα complexes in aged resistance arteries indicate that RSK2 is upstream of mechanisms that regulate the structure, eNOS and Hbα distribution and ultimately NO bioavailabilty. Age related changes in the IEL holes have also been observed previously ^27^. A decrease in Hbα at the MEJ results in a significant decrease in BP in normotensive and hypertensive mice ^15^. Thus, if its inhibitory association with eNOS is decreased in the aged *Rsk2*^*-/-*^ mice MEJs, in conjunction with the normal distribution of eNOS, this would favor NO production and prorelaxant vasodilation and their observed lower BP. Our findings support the hypothesis that MEJs play an important role in BP regulation ^15^.

Whether remodeling of the vasculature is the cause or result of hypertension and whether increased vascular stiffness is the cause of hypertension associated with aging are still open questions. We did not observe significant vascular remodeling associated with aging in arteries from 18-20 month old WT or *Rsk2*^*-/-*^ mice, also reported at 12 months age ^5^ as was a dissociation of vessel stiffness/remodeling and BP ^5,7^. Unlike these studies, in our study mesenteric artery stiffness was increased in both the aged *Rsk2*^*+/+*^ and the aged *Rsk2*^*-/-*^ in the absence of remodeling, suggesting that the increased stiffness develops in late-stage aging and is independent of BP. Thus, the driver for increased stiffness may not be the increased BP in these animals. Alternatively, increased BP could be the driver for increased vessel stiffness but normal eNOS/NO vasodilatory pathways in the *Rsk2*^*-/-*^ as suggested by our data, counteract the effects of increased stiffness to maintain normal BP. Advanced glycation end products (AGEs) that crosslink and stiffen collagen and associated with aging ^28^ were increased in both aged *Rsk2*^*+/+*^ and aged *Rsk2*^*-/-*^ compared to young vessels, accounting for our findings increased vessel stiffness.

Overall, we find that in the absence of RSK2, vessel stiffness can be dissociated from the increased BP of aging. We propose that this reflects a down regulation of procontractile and an increase in prorelaxant pathways resulting in normal BP even with stiffer arterioles. We conclude that RSK2 kinase plays a significant role in the increased BP of hypertension associated with aging.

The increased SBP in aged *Rsk2*^*+/+*^ mice could reflect heart failure and cardiac fibrosis with preserved ejection fraction (HFpEF). We found that cardiac ejection fraction and LVM were not different in WT and *Rsk2*^*-/-*^ mice. Only mild interstitial and modest perivascular fibrosis was present in both the aged *Rsk2*^*+/+*^ and *Rsk2*^*-/-*^ hearts compared to young hearts. Thus, the difference in SBP in the aged *Rsk2*^*-/-*^ and *Rsk2*^*+/+*^ mice did not reflect differences in cardiac fibrosis or changes in ejection fraction.

### Perspectives

This study supports our hypothesis that the extra-renal mechanism, RSK2 signaling, directly regulates vascular SM tone contributing to aging-related hypertension and offers potential therapeutic targets. A RSK inhibitor is in clinical trial for metastatic breast cancer ^29^, raising the possibility of repurposing for treatment of hypertension. Future directions should expand to studies in humans and to the investigation of upstream signaling events and the possibility of RSK2 mediated gene regulation. Our novel findings that structure/function of the IEL/MEJ signaling hub is altered in the aged vessel only in WT but not in aged *Rsk2*^*-/-*^ should be investigated to understand whether and how RSK2 acts locally or through modulation of gene expression. As activated RSK2 is known to drive cell motility and phosphorylation of the Rho GEF, LARG and activation of RhoA to increase cell migration and invasion in cancers, its role in the development of atherosclerosis where cell migration plays an important role could be important. Finally, the role of RSK2 in the kidney is an opportunity for investigation.

### Novelty and Relevance

1. What Is New? *Rsk2*^*-/-*^ mice are resistant to the rise in BP associated with aging and to L-NAME-induced hypertension. Resistant artery stiffness increases in *Rsk2*^*-/-*^ mice despite the normal BP, thus dissociating vessel stiffness, often considered the cause of hypertension, from high BP. Increased artery stiffness is not due to vessel wall remodeling or increased collagen content but corelates with increased AGE mediated collagen crosslinking. Aging increased SM cell Ca^2+^ events, altered the structure and localization of signaling molecules regulating vasodilation at the IEL/MEJ.
2. What Is Relevant? RSK2 is a significant new signaling mechanism mediating age related hypertension.
3. Clinical/Pathophysiological Implications? Our findings present potential new therapeutic targets for treatment of hypertension.

## Nonstandard Abbreviations and Acronyms

cGMP: cyclic guanosine monophosphate
EC: endothelial cell
eNOS: endothelial nitric oxide synthase
Hbα: hemoglobin α
IEL: internal elastic lamina
L-NAME: N(G)-Nitro-L-arginine methyl ester
L-NNA: N^G^-nitro-L-arginine
MEJ: myoendothelial junction
MLCK: myosin light chain kinase
NO: nitric oxide
PKG: cyclic GMP-dependent protein kinase
NHE-1: Na^+^/H^+^ exchanger
RLC_20_: 20 kD myosin regulatory light chain
RSK2: p90 ribosomal S6 kinase
SMC: smooth muscle cell

## Acknowledgments

We thank W.J. Leonard (NIH) for providing the *Rsk2*^*-/-*^ strain, Dr. B.L. Roth Univ of North Carolina for generation of the targeting vector for the RSK2 floxed mice and Taconic Farms for generation of the floxed mice. We thank Dr. Mykhaylo Artamonov, Walter Reed Army Institute of Research, Silver Spring MD., for consultation on assays. We are grateful to the National Institute for Aging at the NIH for providing aged *Rsk2*^*+/+*^ mice to supplement our colony. Figure 6 is created with www.BioRender.com

## Supplementary Figures

### Supplementary Figure Legends

**Figure 1.**
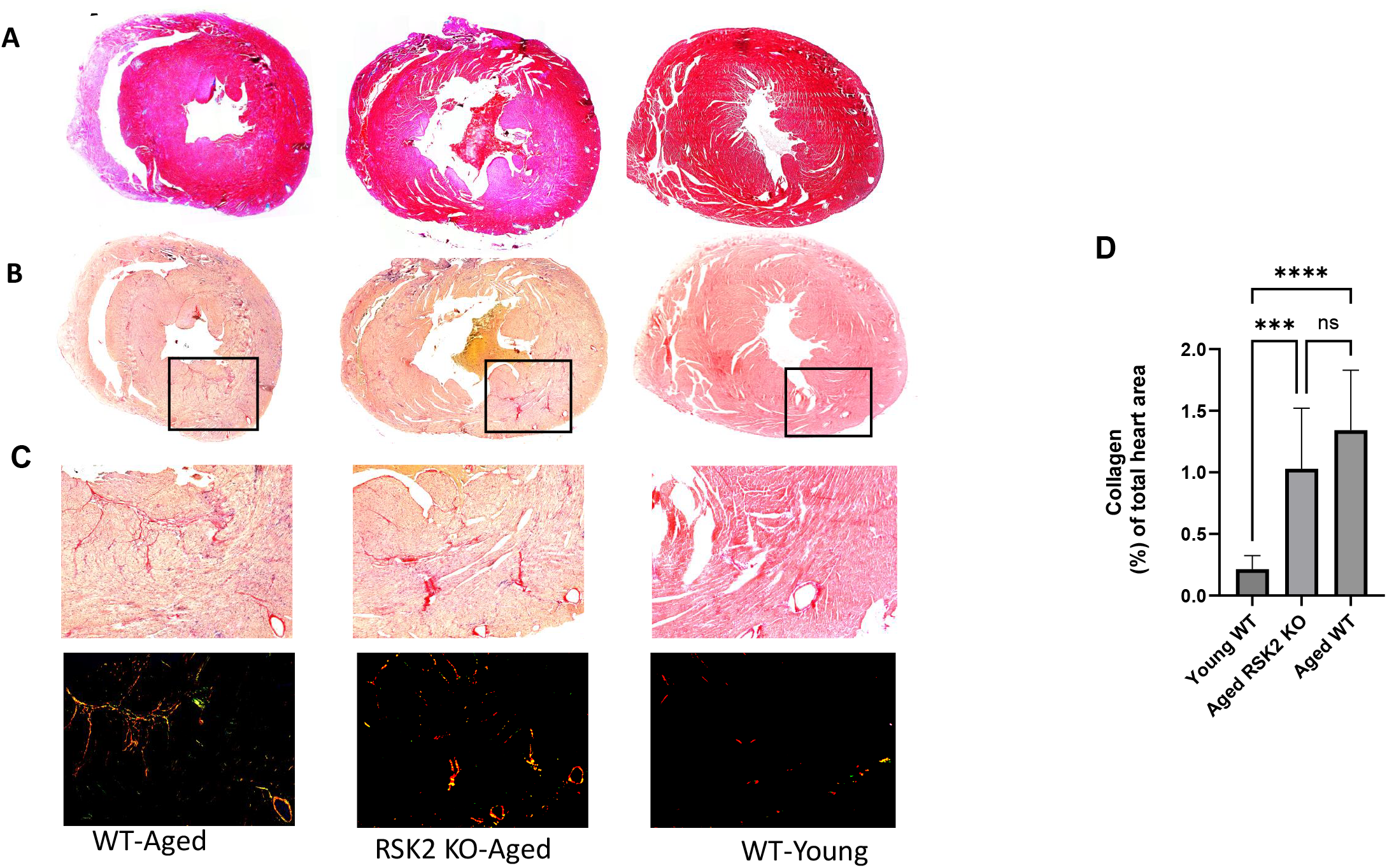
Effect of age and RSK2 expression on extracellular collagen deposition in the heart. Masson’s trichrome (**A**), Picrosirius red staining (**B, C**) and Picrosirius red staining imaged with polarized light (**D**) represent heart cross-sections of aged-wild type (WT), aged-RSK2 deficient (KO) or control young-WT mice. Scale bar 100 µm. (**E**) Quantification of collagen from **D** expressed as a (%) of red fluorescence relative to the total heart area, n= biological samples/condition. Values are presented as means ± SD.

## Supplemental Material

### Methods

#### Animals

All animal studies were performed under protocols that comply with the Guide for the Care and Use of Laboratory Animals (NIH publication no. 85-23, Revised 1996) and were approved by the Animal Care and Use Committee at our institution.

#### *Rsk2*^*-/-*^ mice

*Rsk2*^*-/-*^ mice On a C57BL6 background were a gift from Dr. Warren J. Leonard, National Institutes of Health^30^: they were generated as reported previously ^31^. Aged animals used in this study were 18-20 months old.

#### *Rsk2*_*SM*_^*-/-*^ (*Rsk2Myh11-Cre*^*+*^)mice

The targeting vector was generated by Dr. Bryan Roth’s laboratory, Univ. North Carolina ^32^ and subsequently transfected into the TaconicArtemis C57BL/6N Tac ES cell line by Taconic Farms. Briefly, the RSK2 targeting vector had RSK2 exon 4 flanked by loxP sites, and two positive selection markers flanked by flippase (Flp) recognition target sites (NeoR, neoresistance) and F3 (PuroR, Puromycin resistance) were inserted into introns 3 and 4 respectively (31). Taconic Farms injected the targeted ES cells into blastocysts to generate chimeras. Chimerism was measured by coat color and highly chimeric mice were bred with Flp-Deleter mice to remove selection markers^32^. Sperm from these mice was cryopreserved and for our needs was revitalized by in vitro fertilization by Taconic Biosciences GmbH and its affiliate Taconic Biosciences Inc. using oocytes from superovulated C57BL/6NT ac females. 2 cell embryos were transplanted into recipient females and germline transmission identified in offspring. PCR primers for identification of *Rsk2*_*SM-/-*_ and *Rsk2*_*SM+/+*_ allelles (*Rsk2*^*fl/fl*^) were:

Forward: CCTTAAGTTACAACCCTAGCATCC

Reverse: GTGGAATGTGGATGCTGAGC

Control

Forward: GAGACTCTGGCTACTCATCC

Reverse: CCTTCAGCAAGAGCTGGGGAC

The *Rsk2* ^*flox/-*^ mouse was crossed with a SM myosin heavy chain CreER^T2^ mouse to generate RSK2 knockdown specifically in SM. SM-specific knockout of RSK2 was induced by feeding tamoxifen (40 mg kg^−1^ day^−1^, Envigo Diet TD.130856) to 6-week-old *RSK2*^*fl/fl*^ *Myh11-Cre* mice for 14 days. Mice were used for experiments after a 1-week tamoxifen washout period.

#### Pressure myography

Freshly isolated 3^rd^ or 4^th^ order mesenteric arteries (<180 µm) were placed into HEPES-Krebs solution (mM): NaCl 118.4, KCl 4.7, MgSO_4_ 1.2, NaHCO_3_ 4, KH_2_PO_4_ 1.2, CaCl_2_ 2, HEPES 10, glucose 6, then mounted in a pressure arteriograph (Danish MyoTechnology) as previously described ^33^. The vessels were pressurized to 20 mm Hg and then to 60 or 80 mm Hg, at 37^°^C and equilibrated for 30 min, brought back to 20 mm Hg and then exposed to incremental increases in pressure to 100 mm Hg. Vessels were used only if they displayed robust endothelial cell viability which was assessed at the end of an agonist-induced constriction using 10 µM acetylcholine (Fig. S1C)^33^ or the IK or SK agonist NS309 to induce relaxation of a myogenic constriction ^34^. Luminal diameters were measured in response to changes in intraluminal pressure or to cumulative concentrations of activators or inhibitors applied to the circulating bath. To examine whether the sensitivity to NO is altered in aged mice, exogenous NO donor spermine NONOate (0.3-30 μM) was added to pressurized (80 mmHg) mesenteric arteries from young and aged mice preconstricted with 100 µM Nω-nitro-L-arginine (L-NNA) and the % relaxation determined. Maximum inner diameter was measured after washing with a Ca^2+^ free Krebs-HEPES solution supplemented with 1 mM ethyleneglycol-O, O′-bis(2-aminoethyl)-N,N,N′,N′-tetraacetic acid (EGTA) and 10 µM sodium nitroprusside. Quantification of vessel diameter was performed using the DMT software MyoView. Basal tone and vasoconstriction values were calculated as: [maximum diameter – active diameter / maximum diameter] x 100. Responses are expressed as the percentage of the maximum inner diameter.

#### Measurement of Blood Pressure

Systolic blood pressure was measured in conscious male and female mice (age 26-29 month, *Rsk2*^*+/+*^ mice and their WT littermates) by tail cuff using a MC4000MSP system (Hatteras Instruments, Inc.). Tail cuff was used as these old animals were frail and survival from telemetry surgery was poor. Animals were conditioned by placing them on the apparatus platform for 15 min/day, on 2 consecutive weeks and “sham” measurements were taken. On the next 2 consecutive weeks, mice were placed on the platform daily and at least 10 readings were taken. Conditioning and all blood pressure readings were performed at the same location, by the same operator, at the same time of day under quiet, low-light conditions. Radiotelemetric blood pressure monitoring was used in the young (4-5 month old mice) WT and *Rsk2Myh11-Cre*^*+*^conscious male mice of corresponding age under unrestrained conditions. Continuous blood pressure measurements were performed using Dataquest A.R.T. 20 software (Data Sciences International, St. Paul, MN), as described previously ^35^. Mice were allowed to recover for 7 days after surgery to regain their normal circadian rhythms before blood pressure measurements were initiated. The values over 7-24 hour period were averaged to obtain the baseline day-or night-time blood pressures. Hypertension was induced by administration of L-NAME in the drinking water at a concentration of 0.5 mg/mL and changed 3 times/week, for a targeted dose of 40-60 mg/kg/day. Measurement of BP under L-NAME were taken during the 2^nd^ week of L-NAME exposure.

#### Cardiac Physiology Measurements

Echocardiography was performed using a Vevo3100 ultrasound (FUJIFILM VisualSonics, Toronto, Canada) and a linear-array 40MHz transducer (MS-550D). Left ventricle systolic and diastolic measurements were captured in M-mode from the parasternal short axis.

#### Western blotting

Abdominal aorta and mesenteric artery arcades were used for detection of total proteins and phosphorylation of RSK2^Ser227^. Arteries were equilibrated at in HEPES-Krebs solution at 37°> C for 30 mins prior to snap freezing in liquid N2, transfer to vials of 10% trichloroacetic acid in acetone in a liquid N2 atmosphere and stored at -80°C for freeze substitution, as described previously ^36^. Freeze substituted arteries were washed 3X in acetone, dried and homogenized in glass-on-glass hand-operated micro homogenizers, in 2X Laemmli Sample Buffer and urea. Samples where heated to 100^°^C for 5 min and subjected electrophoresis and western blotting.

#### Histology

After fixation of iliac and mesenteric arteries and hearts in 10% buffered formalin, alcohol dehydration and paraffin embedding, 4-5 μm sections of paraffin-embedded blood vessels or 7 μm thick sections of heart tissues were processed and stained with Picrosirus red and Masson’s trichrome for analysis of collagen content and for measurements of vessel wall and lumen. Images were captured with brightfield and with polarized light for the Picrosirus red staining, according to standard procedures.

#### Immunohistochemistry for detection of advanced glycation end products (AGEs)

Antigen retrieval was performed on paraffin embedded iliac artery sections (5μm thick) with TE buffer pH8.0 in a pressure cook for 15 minutes, followed by 1% H2O2 to block endogenous peroxidase. After blocking with 10% goat serum, tissue sections were incubated with mouse monoclonal anti-AGE (1:50) antibody overnight at 4 ^°^C. Biotinylated goat anti-mouse IgG (1:200, Vector Laboratories) was applied for 1 hour, followed by avidin and biotinylated enzyme. The signal was visualized with 3′-diaminobenzidine (DAB). Hematoxylin was used for counterstaining. IgG control was performed following identical procedures excluding the primary antibody incubation. Intensity of AGE product was scored by a blinded investigator.

#### Quantification of fibrosis using polarized light microscopy image analysis

The Picrosirius red stained heart sections were analyzed under polarized light using a Leica Thunder Imager 3D Cell Culture with Leica DMD108 digital microscope fitted with polarization filters to allow only polarized light to pass through and be transferred into a binary image, used for the calculation of fibrosis. Images were obtained with a 10x objective and a monochrome charge-coupled device camera and analyzed using ImageJ software. A threshold was set to show areas only occupied by collagen. Subsequently, binary images were used to calculate the fibrotic area as the percentage of collagen per the total tissue area.

#### Ca^2+^ measurements

Ca^2+^ signals in third-order branches of mesenteric arteries were imaged using an Andor Revolution WD (with Borealis) spinning disk confocal imaging system (Andor Technology, Belfast, UK), comprised of an upright Nikon microscope with a 40X water dipping objective (numerical aperture 0.8) and an electron multiplying charge coupled device camera^8,34^ Pressurized (80mm Hg) third-order branches of mesenteric arteries (∼100 μm internal diameter at 80 mmHg) were loaded with fluo-4 AM (10 μM) for 1 hr, to measure calcium signaling events in the mesenteric artery SMCs of young (13 weeks), aged (88 weeks), prior and following addition of the RSK2 inhibitor, LJH685 (5μM). Ca^2+^ frequencies (Hz) and amplitudes (F/Fo) were measured under the above conditions Images were recorded at 60 frames s−1. Fluo4-bound Ca^2+^ was detected by exciting at 488 nm with a solid-state laser and collecting emitted fluorescence using a 527.5–49 nm bandpass filter.

#### Vessel Stiffness

3^rd^ or 4^th^ order mesenteric arteries from young and aged *Rsk2*^*+/+*^ and *Rsk2*^*-/-*^ mice were mounted on a pressure myograph and studied under Ca^2+^-free conditions to eliminate the contractile mechanisms and fully relax the vessels. Arterial stiffness was calculated from passive-pressure diameters taken at 10, 20, 40, 60, 80, 100, and 120 mmHg. The wall:lumen ratio was determined in the young and aged *Rsk2*^*+/+*^ and *Rsk2*^*-/-*^ mice. Assessment of arterial stiffness was determined by comparing the rate constant of an exponential function fitted to the stress strain curve for each vessel^37,38^. Stress-strain curves were calculated: circumferential stress = (intraluminal pressure x lumen diameter)/(2 x wall thickness); strain = (D-D0)/D0, where D is the inner diameter at a given pressure step and D0 is the inner diameter at 10 mmHg.

#### *In situ* proximity ligation assay (PLA)

Mouse second-order MAs were isolated and pinned down *en face* on a SYLGARD block. Arteries were fixed in PFA (4%) for 10 minutes, washed with PBS and permeabilized by incubating in a solution containing 0.2% Triton X-100 for 30 minutes at room temperature. Arteries were incubated with a blocking solution containing 5% normal donkey serum for 1 hour at room temperature. Samples were incubated overnight at 4C with anti-eNOS (1:500, 610297, BD Bioscience) and anti-Hemoglobin α (1:500, ab92492, Abcam) antibodies. The following day the protocol for the Duolink PLA Technology kit (MilliporeSigma) was followed according to the manufacturer’s instructions. Nuclei were stained using 0.3 μM DAPI (Invitrogen). Images were acquired using the Zeiss LSM 700 confocal microscope. Images were captured along the z-axis at a slice thickness of 0.2 µm. Image analysis was performed using ImageJ (NIH). Images were normalized by the number of positive puncta by the number of nuclei in the field of view.

#### Antibodies

Proteins were transferred to polyvinylidene difluoride membranes and blocked with Odyssey Blocking Buffer, probed with primary antibodies in Blocking Buffer and detected and quantified on the Odyssey system (Li-Cor). Li-Cor Total Protein stain was used for loading normalization. The following antibodies were used: mouse monoclonal and rabbit polyclonal anti-actin (1:5,000 western blot (WB); Sigma-Aldrich); rabbit polyclonal antibodies anti-RSK2 phospho-Ser^227^ and mouse monoclonal anti-RSK2 (1:500 WB; 1:50 IP, Santa Cruz Biotechnology Inc.); mouse monoclonal anti-AGE (1:50) antibody (MyBioSource); GAPDH (1:5,000 WB; Millipore). Primary antibodies were detected using either goat anti-rabbit and anti-mouse Alexa 680 (1:10,000, Invitrogen), donkey anti-sheep Alexa 680 (1:10,000, Invitrogen), or a goat anti-rabbit and anti-mouse IRDye800 (1:10,000, Rockland Immunochemicals)-conjugated secondary antibody. Representative western blots have been cropped for presentation.

#### Data Analysis

All statistical analyses were performed using GraphPad Prism (Version 10.4.0). Data are presented as the means ± S.E.M. The normality of data distribution was assessed using the D’Agostino & Pearson test. A two-tailed unpaired t-test with Welch’s correction was used, for comparisons between two groups. Brown-Forsythe and Welch ANOVA was performed, followed by a Dunnett T3 test for comparisons among multiple groups. The level of significance was set at P< 0.05.

